# Ghrelin inhibits courtship ultrasonic vocalization in male mice

**DOI:** 10.64898/2026.09.21.753339

**Authors:** Rachel G. Bond, Luke T. Brown, Max Otto Jones, Joel A. Tripp

## Abstract

Courtship is an essential component of reproductive success for many species and many courtship behaviors, such as vocalization, are energetically demanding. In order to successfully balance investment in courtship with feeding and foraging, individuals must be aware of their current energy balance, and hormonal signals of energetic state have been found to influence social signal effort across a variety of species. Ghrelin is one hormone that plays an important role in regulating energy balance by signaling hunger and promoting feeding. Previous studies have investigated the role of ghrelin in courtship, aggression, and mating behavior. In this study, we build on prior work by investigating the effects of ghrelin on ultrasonic vocalizations (USV) and other courtship behaviors of male mice. We expand on previous research by observing ghrelin’s effects over longer interactions, allowing us to understand how ghrelin impacts the dynamics of USV production over time. Our results show that while males initially vocally court females at a high rate regardless of treatment, ghrelin reduces overall USV production by increasing the rate of decline of vocalization. Surprisingly, despite its effects on vocal courtship, we found no significant differences in the production or timing of other, non-vocal, courtship behaviors. This study deepens our understanding of the role of hormones traditionally associated with controlling hunger and feeding in regulating social behaviors.

## Introduction

Successful reproduction depends on an animal’s ability to first locate, attract, and court a mate. Courtship displays are ritualized sets of signaling behaviors that serve to stimulate and synchronize mating as well as convey signaler quality to the intended mate (Mitoyen et al., 2019). Vocalization is a common component of courtship display that is often energetically costly. Calling is estimated to increase energy expenditure by eight-fold in insects and amphibians and doubles metabolic rate over rest in birds (Ophir et al., 2010). Although the direct energetic cost of vocalization may be smaller, but still significant, in mammals (Chaverri et al., 2021; Collier et al., 2022; Ilany et al., 2013; Noren et al., 2013), callers also face indirect costs such as the potential to attract a predator (Mougeot and Bretagnolle, 2000; Tuttle and Ryan, 1981) or competitor (Hammond and Bailey, 2003) and the loss of foraging opportunities (Abrahams, 1993; Kim et al., 2008).

Despite these potential costs, energy investment in courtship displays has been consistently linked to greater reproductive success across the animal kingdom (Mitoyen et al., 2019). Thus, an individual’s capacity to invest in courting is inherently tied to that individual’s internal energy state. Additionally, hormonal signals of energy balance have been shown to directly regulate the effort animals invest in social displays (Giglio and Phelps, 2020; Sinnett and Markham, 2015; Tripp et al., 2026). Elucidating how internal energy homeostasis regulates an animal’s motivation and ability to invest in courtship is therefore necessary for understanding the conditions under which reproduction succeeds.

Because it is metabolically demanding to produce courtship displays and engage in other appetitive mating behaviors, hormones and neuromodulators that serve as signals for an animal’s internal energy state are in a strong position to regulate investment in courting a mate. One such hormone is ghrelin, an orexigenic peptide primarily responsible for signaling hunger and promoting feeding behavior (Cummings et al., 2001). Ghrelin acts on growth hormone secretagogue receptor 1A, a receptor with a known role in regulating metabolism and stimulating appetite and growth hormone release (Yin et al., 2014). Roughly 60-70% of circulating ghrelin is produced in the stomach, and the rest produced in tissues such as the small intestine, pituitary, hypothalamus, and pancreas (Tritos and Kokkotou, 2006). Beyond its role in signaling hunger, ghrelin has been shown to influence cardiovascular function (Korbonits et al., 2004), central reward and motivational circuits (Engel and Jerlhag, 2014), reproductive function (Tena-Sempere, 2007), and aggression (Shah and Nyby, 2010). However, studies investigating the role of ghrelin in courtship and mating have yielded conflicting results.

Male mice have been found to produce significantly fewer USVs in brief interactions following peripheral ghrelin injection, which suggests that they reduce courtship investment when ghrelin levels are high (Shah and Nyby, 2010). In contrast, engagement in and motivation toward sexual behaviors, namely mounting, has been found to significantly increase in male mice receiving similar peripheral ghrelin treatment (Egecioglu et al., 2016), and ghrelin signaling in the ventral tegmental area plays an important role in sexual motivation in rats (Hyland et al. 2018). In contrast, central ghrelin administration in male rats increases the latency to mount and the number of mounts required before ejaculation, while decreasing overall copulation success (Babaei-Balderlou et al. 2016). Collectively, these findings suggest that the effects of ghrelin on courtship and mating depend on the species and strain tested as well as the location, dose, and duration of administration, which have not been systematically compared within a single model.

House mice (*Mus musculus*) provide a useful model for studying how internal energy states influence an animal’s ability to invest in courtship displays. Male mice court females using ultrasonic vocalizations (USVs) alongside behaviors like sniffing and mounting, and females are attracted to male USVs (Egnor and Seagraves, 2016). Courtship USVs in mice are surprisingly complex, composed of multiple distinct syllable types that are organized into discrete, structured sequences (Klaus et al., 2025). Males producing calls with longer syllable durations tend to produce vocalizations at a higher rate, and males with more complex vocal bouts have been observed to achieve greater copulatory success (Kanno and Kikusui, 2018; Klaus et al., 2025; Nicolakis et al., 2020).

In this study, we sought to build on previous experiments by examining the effects of acute ghrelin administration on vocal courtship behavior over a longer time period than previously observed. This allowed us to characterize the effects of ghrelin on the dynamics of courtship USV production. In addition to USVs, we also quantified non-vocal social behaviors related to courtship and mating. We hypothesized that exogenous peripheral ghrelin would signal a lower energetic state that would reduce male investment in courtship, resulting in fewer USVs and non-vocal courtship behaviors. Surprisingly, we found that while ghrelin did reduce vocal courtship behavior, the hormone had little effect on non-vocal behaviors during courtship interactions.

## Materials and Methods

### Animal Subjects

Subjects were adult (11-16 weeks old) sexually naive male C57BL/6J (Strain #000664) mice obtained from The Jackson Laboratory (Bar Harbor, ME) at 8 weeks of age. Four adult (12-17 weeks old) female C57BL/6J mice were used as stimuli. Stimulus females were ovariectomized to prevent pregnancy and avoid potential effects of estrous cycle on stimulus animal behavior. Ovariectomies were performed under ketamine (50-80mkg/kg) and xylazine (5 mg/kg) anesthesia. Briefly, a small incision was made in the abdomen, the uterine horns and ovaries were gently separated from the periovarian fat pads, each uterine horn was tied with a suture, the ovaries were removed, abdominal wall sutured, and incision closed with a surgical staple. Females were sexually naive prior to the experiment and were used as stimuli five times each, with at least one week to recover between trials. Male mice were housed in groups of four in 19 in. x 10.5 in x 6.125 in. acrylic cages (Ancare, Bellmore, NY) with a mouse igloo and *ad libitum* access to water and rodent chow (Teklad Rodent Diet 2018, Inotiv, West Lafayette, IN). Female stimulus mice were housed in pairs in cages of the same dimensions and conditions. Animals were held on a 12:12 light:dark cycle. All procedures followed the ARRIVE guidelines and were approved by the Institutional Animal Care and Use Committee of Carleton College.

### Drugs

We dissolved ghrelin (R&D Systems, Minneapolis, MN) in sterile saline at 0.033 mg/mL and stored at −20℃ for up to one month. Before injecting in subjects, we thawed the solution completely and allowed it to reach room temperature. Ghrelin dosage was based on prior studies of ghrelin effects on social behavior (Egecioglu et al., 2016; Shah and Nyby, 2010).

### Experiment Design

We randomly assigned each male subject to receive either ghrelin or saline injection. We weighed and injected each subject intraperitoneally with ghrelin (0.33 mg/kg at 0.033 mg/mL in sterile saline) or an equivalent volume of sterile saline. We placed each male in an empty 19 in. x 10.5 in x 6.125 in. cage with bedding to acclimate for 20 minutes. After acclimating, we placed a stimulus female into the center of the cage. The male and female were video recorded using a high-definition video camera (Vivitar DVR8K-BLK-STK-4, Edison, NJ) and audio was recorded using a Dodotronic UltraMic 250K microphone (Castel Gandolfo, Italy) placed directly above cages and Audacity software at a sample rate of 250000 hZ. After 30 minutes, we ended the video and audio recording and returned mice to their home cages. Two males, one ghrelin-treated and one saline-treated, were tested in the same room simultaneously per trial. Testing occurred during the animals’ light phase, and we conducted half of the trials in the morning between one and three hours after light onset (08:00-10:00) and half in the afternoon between six and eight hours after light onset (13:00-15:00).

**Figure 1.**
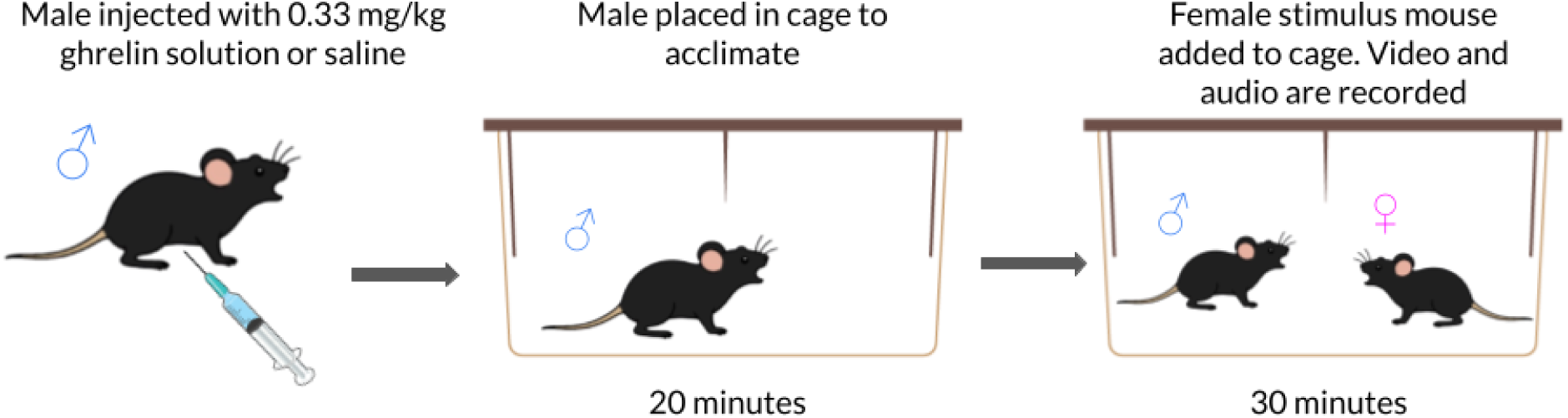
Experiment design. Male mice were weighed, injected with ghrelin or equivalent volume of saline, then acclimated in an empty cage for 20 minutes. An ovariectomized female was added to the cage, and behaviors were recorded for 30 minutes. Figure created using illustrations from NIAID NIH BioArt Source (NIAID Visual & Medical Arts, 2024) and SciDraw (Li, 2026).

### Behavioral Analyses

We used DeepSqueak (Coffey et al., 2019) to identify and analyze USVs. Calls between 40-125 kHz were detected using the Mouse Detector YOLO R2 network and detections were verified by a trained human observer. We compared the number of USVs and length of individual calls as determined by DeepSqueak analysis. Similar to previous studies (Castellucci et al., 2018; Marconi et al., 2020), we defined USV bouts as a series of syllables with an inter-call interval of less than 300 ms. The data from two mice per treatment were not included for our analyses of vocal behavior as their corresponding audio files did not contain the full 30-minute trial.

To analyze non-vocal behaviors, video footage of each trial was imported into BORIS (Friard and Gamba, 2016). We quantified four social behaviors: durations of following, social sniffing, and anogenital investigation (AGI), and the number of mounting attempts. Following was defined as the male pursuing behind the female within one tail length. Social sniffing was defined as a face-to-face investigation or sniffing around the body of the female while not engaged in AGI or following. AGI was defined as the male putting its nose in the anogenital area of the female, either under the tail from behind or under the leg from side of the female. Mounting was defined as the male placing forepaws on the female haunch and attempting intromission. We quantified the duration of following, social sniffing, and AGI as well as the number of mounting attempts and the latency for each behavior. In addition to these social behaviors, we also quantified the time subjects spent roaming the cage, which included walking, rearing, or climbing in the absence of social interaction and the time spent stationary, in which the subject was immobile with all paws on the cage floor while not engaged in other identifiable behaviors (such as grooming) or social interaction.

Each behavior was scored by two researchers blinded to the subject’s condition. To assess consistency between observers, we calculated the Pearson correlation for the total duration or occurrences quantified by the two observers for each behavior. We found a significant (p < 0.0001) positive correlation for each behavior of interest, with an average correlation coefficient of 0.923 (range: 0.817 for social sniffing to 0.974 for roaming). We used the average values for duration, occurrences, and latency across the two observers for our statistical analysis.

### Statistical analysis

All analyses were conducted using R (v4.6.0) in the RStudio (v2026.08.1+195) environment or the Julia Programming Language (v1.12.2). For each comparison, we checked whether the data were normally distributed using the Shapiro-Wilk test. For data that did not significantly differ from normality, we compared across treatment groups using a Welch Two-Sample t-test. These included USV counts, call length, number of USV bouts, bout length, peak USV rate, and durations of following, social sniffing, AGI, and roaming. When our data significantly varied (p < 0.05) from a normal distribution, we instead used the Wilcoxon rank sum exact test. These included mounting occurrences, stationary duration, and latencies to following, social sniffing, AGI, and mounting.

To assess the relationship between USV rate and call duration we calculated a Pearson’s correlation coefficient separately for the ghrelin- and saline-treated groups and for both groups together. To assess the time course of vocalization, we compared USV production in 90 second bins, aligned to each subject’s first call, using a generalized linear mixed model with treatment, time, and treatment × time interaction as fixed effects and subject and observation as random effects. Because most mice began calling at high rates with short latencies, this approach best allowed us to describe calling dynamics at a group level. Effect sizes were calculated using Psychometrica (https://www.psychometrica.de/effect_size.html) and are reported as Cohen’s *d* for all pairwise comparisons, as rate ratios with Wald 95% confidence intervals (CI) for the generalized linear mixed model, and as model-implied percent decline in expected USV count per minute with delta-method 95% confidence intervals.

## Results

### Ghrelin reduces courtship USV production

To determine ghrelin’s overall impact on vocal courtship behavior, we first examined the effect of ghrelin treatment on USV production and call characteristics (Figure 2). Males with elevated ghrelin levels produced significantly fewer USVs than control males (t_11.68_ = −2.27, p = 0.043, *d* = 1.13). On average, control males produced more than double the number of USVs of ghrelin-treated males. However, we found that ghrelin treatment did not affect the length of calls that were produced (t_14_ = −0.566, p = 0.58, *d* = 0.28).

**Figure 2.**
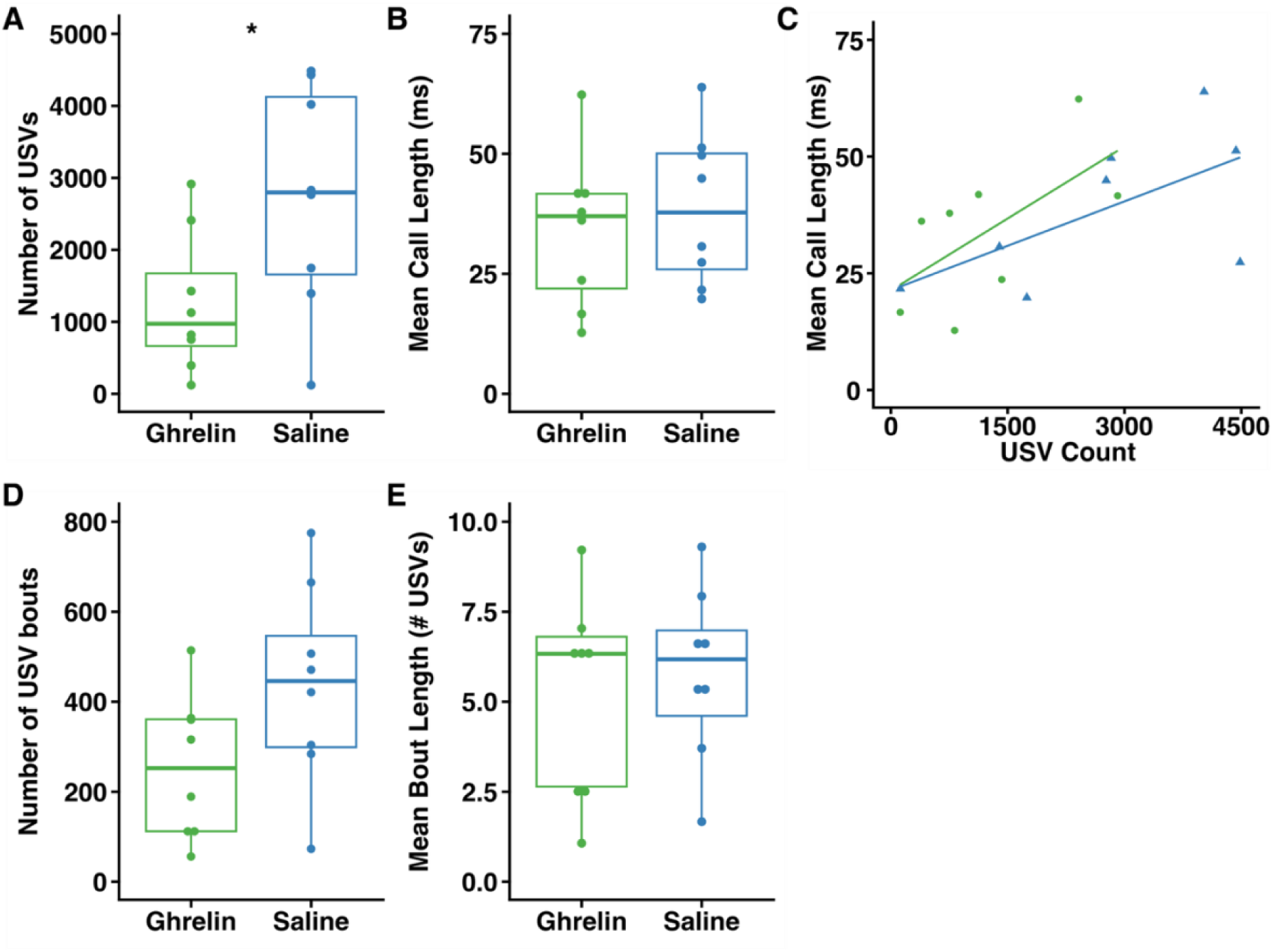
Ghrelin reduces courtship USV production. A) Ghrelin-treated male mice (n = 8) produced fewer USVs during a courtship assay compared to saline-treated control (n=8) mice but produced calls of similar length (B). C) Call length was positively associated with the number of USVs produced. We found no significant effect of ghrelin on D) number of USV bouts, or E) mean length of bouts. Boxes show first and third quartiles, bar indicates median, and whiskers extend to maximum and minimum values. Points show data from individual mice. * denotes statistically significant difference (p < 0.05).

Despite this, we found a positive relationship between the number of USVs produced and mean call length for both the ghrelin (r = 0.620) and saline (r = 0.621) groups. While this relationship was not statistically significant when comparing groups separately (p = 0.10 for each comparison), when both groups were considered together, we found a similar positive relationship that was statistically significant (r = 0.59, p = 0.016) Additionally, while ghrelin-treated males tended to produce slightly fewer USV bouts neither the number of bouts nor the length of bouts differed significantly across groups (bouts: t_12.70_ = −1.91, p = 0.079, *d* = 0.96; bout length: t_13.63_ = −0.46, p = 0.65, *d* = 0.23).

### Ghrelin impairs the persistence of, not the capacity for, male courtship vocalization

Although ghrelin-treated males produced significantly fewer total USVs (Figure 2A), this measure alone cannot distinguish whether ghrelin reduces calling rate throughout courtship interactions, resulting in reduced peak USV rates, or if ghrelin treatment results from faster rate decay from a similar peak (Figure 3A). To dissociate these possibilities, we examined USV activity across the session for individual subjects and as a group average (Figure 3B-D). Raster plots of individual mice showed that while both groups typically produced USVs at high rates at the beginning of trials, control (saline-treated) males were more likely to sustain frequent, consistent USV production throughout the 30-minute assay, while ghrelin-treated males’ USVs were concentrated earlier in the session and dropped off for most individuals as the assay continued.

**Figure 3.**
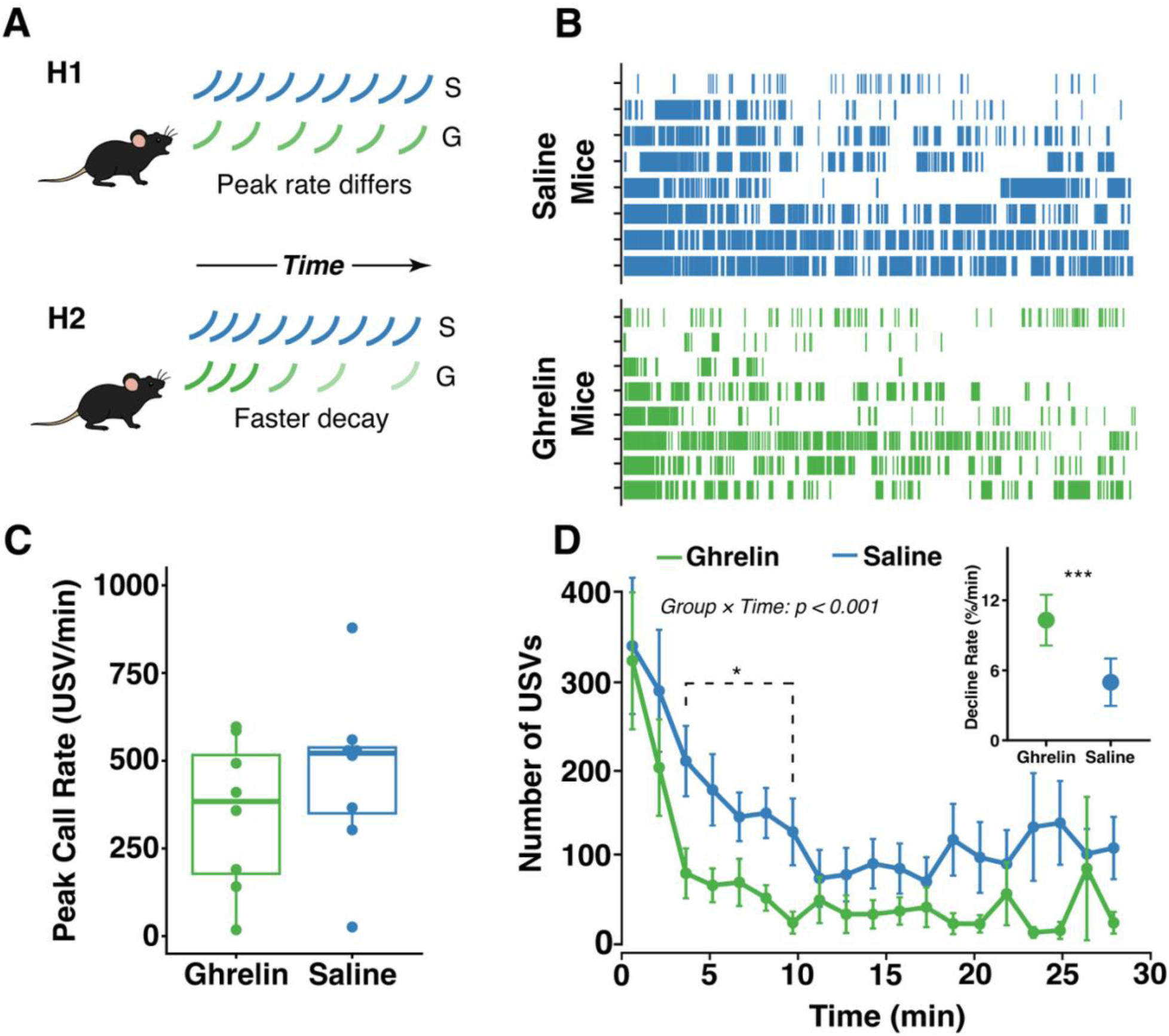
Ghrelin influences the dynamics of male courtship vocalization. **A)** USV rate could differ due to differences in peak call rate and rate throughout the trial (H1) or due to faster decay from peak calling rate at the start of the trial (H2). **B)** Raster plots of USVs produced by saline- and ghrelin-treated mice. Each tick represents one call, each line shows 30-minute trial for an individual mouse. **C)** Peak USV rate does not differ by hormone treatment. Boxes show first and third quartiles, bar indicates median, and whiskers extend to maximum and minimum values. Points show data from individual mice. **D)** Average USV rate over time. Dots are group average call rates (90 second time bins). Error bars represent standard error of the mean. The dashed bracket and asterisk mark the cluster of bins (3-7, 3.0-10.5 min) in which saline-treated males produced significantly more USVs (cluster-based permutation test, cluster p = 0.021). Decline rate (*inset*) was significantly higher for ghrelin-treated mice compared to saline-treated controls. Dots show model-implied group decline rate and error bars show 95% CI.

Consistent with this observation, we found that peak USV rate was similar across hormone treatment (t_13.75_ = −0.99, p = 0.34, d = 0.50). We also found this pattern reflected in the group-averaged USV rate over time: both groups began the assay with a comparably high USV rate, but ghrelin-treated males showed a significantly steeper decline over time, while control males maintained a higher USV rate for more of the session (Figure 2B). This divergence was confirmed by a population-level generalized linear mixed model, which identified a significant treatment × time interaction (β = −0.058 ± 0.016 SE, z = −3.51, p = 0.00045). This interaction corresponds to a rate ratio of 0.944 per minute (95% CI 0.914-0.975): with each additional minute, the ratio of expected ghrelin to saline USV counts fell by a further 5.6%. Model-implied decline rates were 5.0% per minute (95% CI 3.0-7.0) in saline treated males, and more than double that in ghrelin treated males (10.3% per minute, 95% CI 8.1-12.5). Further, to determine when the groups diverged, we compared USV counts at each bin using a cluster-based permutation test. One cluster spanning bins 3-7 (3.0-10.5 min) was significant, with saline treated males producing more USVs (summed t = 12.55, cluster p = 0.021); no other cluster was significant (p > 0.05). Together, these results show that differences in USV rate were not due to a reduced ceiling on USV output, indicating that ghrelin-treated males were capable of producing USVs at rates comparable to saline-injected mice. Instead, the difference was specific to each subject’s likelihood of sustaining that rate over time.

### Ghrelin does not impact non-vocal behaviors

We also examined ghrelin’s effect on non-vocal courtship and social behaviors, but did not find any significant differences between ghrelin-treated and control groups (Figure 4). We first examined the amount of time males spent following, sniffing, and engaging in AGI with females along with the number of mounting attempts made. While ghrelin-treated males spend less time following females on average, this difference was not statistically significant (t_17.38_ = −1.36, p = 0.19, *d* = 0.61). Additionally, we found no differences in the time spent sniffing females (t_17.86_ = −0.0162, p = 0.99, *d* = 0.0070) or engaging in AGI (t_14.68_ = −0.443, p = 0.66, *d* = 0.20). Likewise, there was no significant difference in the number of mounting attempts made across treatment groups (W = 35, p = 0.27, *d* = 0.56). We found no differences in the latencies to engage in any non-vocal social behavior (following: W = 41, p = 0.53, *d* = 0.31; social sniffing: W = 32, p = 0.19, *d* = 0.74; AGI: W = 49, p = 0.97, *d* = 0.23; mounting: W = 64, p = 0.30, *d =* −0.53).

**Figure 4.**
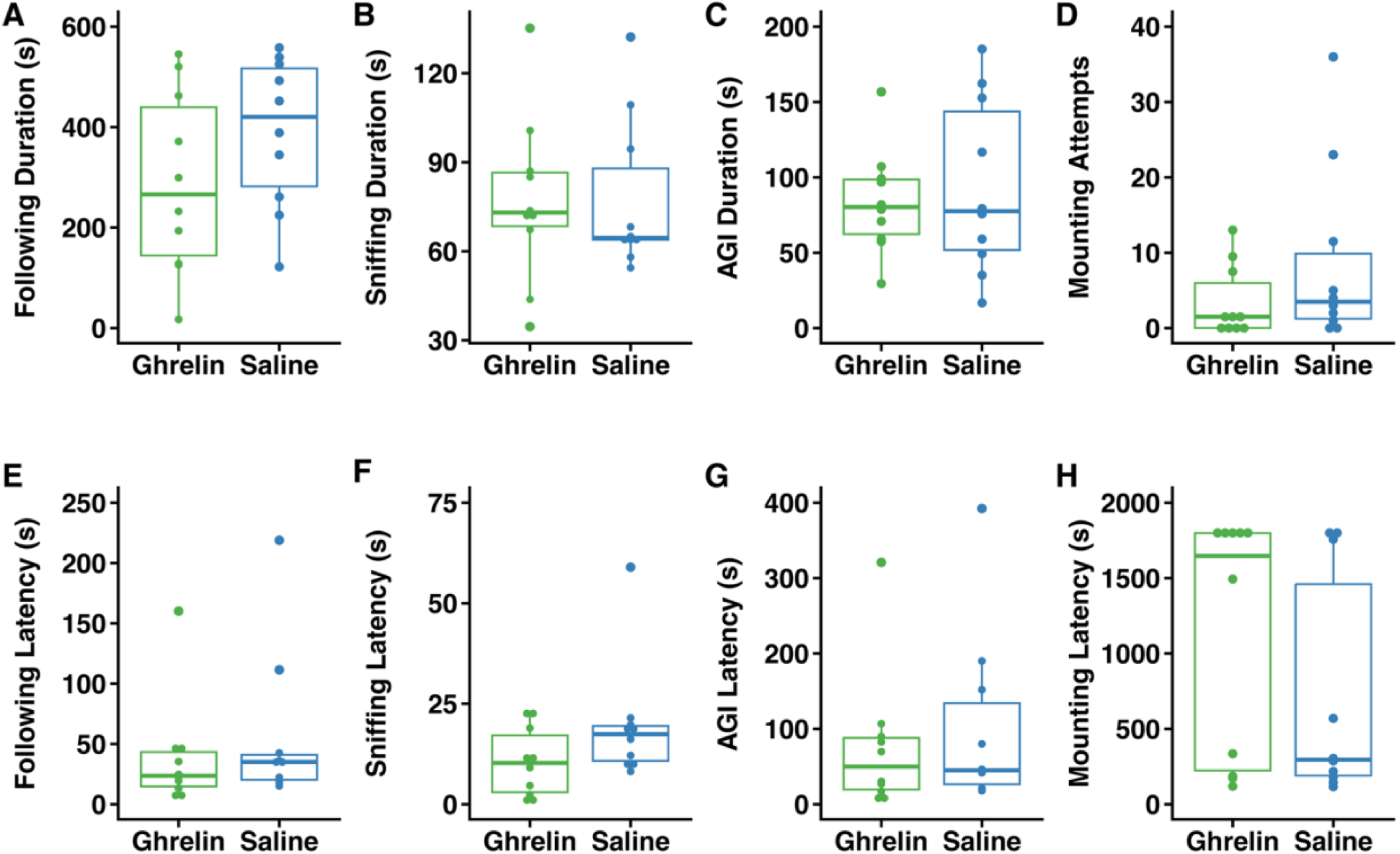
Ghrelin does not influence non-vocal social behaviors. Duration (A-C) or occurrences (D) and latencies (E-H) of non-vocal social behaviors performed by male mice (n = 10 per treatment) during courtship interactions with females, including following (A, E), sniffing (B, F), anogenital investigation (C, G), and mounting (D, H). No significant differences (p > 0.05) found among non-vocal social behaviors. Boxes show first and third quartiles, bar indicates median, and whiskers extend to maximum and minimum values. Points show data from individual mice.

Finally, we assessed whether ghrelin treatment influenced overall levels of activity. We found that neither the time spent roaming the cage (t_17.86_ = 0.056, p = 0.96, *d* = - 0.025) nor time spent stationary (W = 49, p = 0.97, *d* = −0.0040) in the absence of social interaction significantly differed between mice given ghrelin or saline injection.

**Figure 5.**
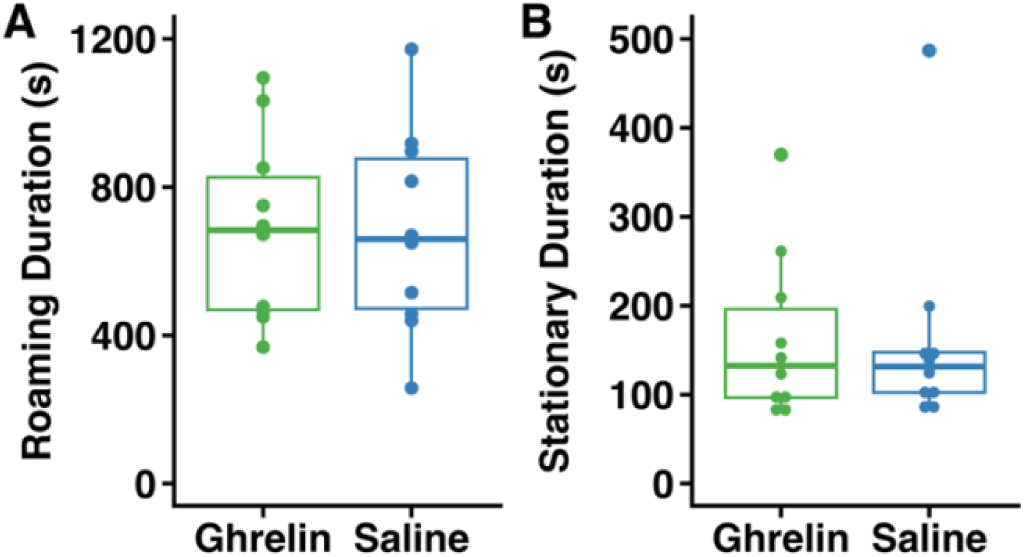
Ghrelin does not influence general activity. Time spent roaming the cage (A) or stationary (B) while not engaged in social interaction did not significantly differ (p > 0.05) across treatment groups (n = 10 per treatment). Boxes show first and third quartiles, bar indicates median, and whiskers extend to maximum and minimum values. Points show data from individual mice.

## Discussion

We found that male mice treated with ghrelin exhibit reduced vocal behavior during courtship with females. Ghrelin-treated male mice displayed fewer USVs, showing a steeper decline in calling rate over the course of the assay. In contrast, none of the four non-vocal courtship behaviors—following, sniffing, anogenital investigation (AGI), and mounting—differed significantly between the groups, whether they were assessed by duration and frequency or latency to first occurrence.

Our main result is consistent with prior work showing that ghrelin reduces USV rate in male-female interactions (Shah and Nyby, 2010). We also extend this finding by looking at a longer time interval for courtship interaction, which allowed us to more deeply investigate the dynamics of USV production and the influence of ghrelin. While mice in both groups produced USVs at high rates at the beginning of trials, resulting in similar peak USV rate, those given ghrelin prior to the trial experienced a larger rate of decline, resulting in fewer USVs produced overall. This difference appears to be largely driven by a reduction in the number of bouts, rather than the number of USVs per bout. While neither of these measures differed significantly, we note that saline-treated mice produced nearly double the number of bouts compared to ghrelin-treated mice, while bout length was similar between groups. Future studies with larger behavioral samples and statistical power may further support this observation.

Beyond vocal courtship, we were quite surprised to find that ghrelin did not significantly influence behaviors related to social interest, courtship, or mating, including following, social sniffing, anogenital investigation, or mounting. The absence of effect on mating-related behaviors was particularly unexpected, as ghrelin has been found to regulate mating in several previous studies; however, the direction of the effect has varied across those studies. For example, ghrelin reduced mounting behavior in male Wistar rats and Swiss mice when injected directly into the brain ventricles or hypothalamus, respectively (Babaei-Balderlou and Khazali, 2016; Belén Poretti et al., 2023). Systemic ghrelin similarly reduced receptivity in ovariectomized and hormonally-primed female Swiss mice (Bertoldi et al., 2011) and injection of ghrelin into the medial preoptic area of Long Evans rats reduced appetitive sex behavior (Hyland et al., 2018). On the other hand, intraperitoneal injection of ghrelin increased preference toward females and mounting in male NMRI mice (Egecioglu et al., 2016), while ghrelin injection into the ventral tegmental area or laterodorsal tegmental area reduced the latency and increased the rate of mounting females (Prieto-Garcia et al., 2015). Taken together, these studies illustrate that the effects of ghrelin on courtship and sexual behavior are complex and may differ depending on the method and target of delivery, species and strain tested, and the specific behavioral assay. Further research on this subject should clarify ghrelin’s role in regulating reproductive behavior by examining its effects through different routes of administration and varied behavioral assays within one model.

Further, we found no effect of ghrelin on general activity levels in our experiment, as measured by time either roaming the cage or remaining stationary in the absence of social interaction. This was somewhat surprising, because ghrelin has been found to increase motivation toward voluntary exercise in the form of wheel running in mice (Mifune et al., 2020; Tezenas Du Montcel et al., 2023); however, in Siberian hamsters this effect was only found when running on a wheel was necessary to gain access to food (Keen-Rhinehart and Bartness, 2005), suggesting that ghrelin-induced activity may be related to foraging behavior. As mice in our study did not have running wheel access and trials took place in relatively small home cages with no access to food, our subjects may not have been motivated to forage. Nevertheless, these results are important in demonstrating that ghrelin treatment had a specific effect on vocal behavior rather than producing a general increase or decrease of activity.

Our goal with this study was to examine how ghrelin’s signaling of hunger influences motivation and investment in courtship behavior. Mating behaviors can be broadly classified into two main categories: appetitive courtship behaviors, which serve as a prerequisite to the “end-goal” of the second category, sexual consummatory behaviors (Ball and Balthazart, 2008). Because appetitive behaviors are a precursor to consummatory ones, measuring courtship behaviors is a critical metric of understanding motivational mechanisms. Although we initially hypothesized that elevated ghrelin would lead to a consistent decrease across both vocal and non-vocal courtship behaviors, our findings suggest that ghrelin’s modulation of courtship most strongly influences investment in courtship USVs.

Courtship vocalization is a social behavior that requires high effort and while it leads to highly-rewarding, consummatory sex behaviors, we found that male mice with elevated ghrelin levels do not persist in vocalizing as long as control mice. We demonstrate that the capacity to call is not inhibited; rather it appears that the motivation to court a female is diminished. A likely explanation for this effect is ghrelin’s impact on internal energy homeostasis. Typically, elevated levels of endogenous ghrelin signals low energy availability by activation of GHSRs and phosphorylation of AMPK in the ventromedial hypothalamus, responsible for upregulating fatty acid synthesis and fat storage (Sovetkina et al., 2020). This pathway mainly stimulates the sensation of hunger but broadly leads to the organism regulating its energy expenditure. As the organism’s internal energy level decreases, they are perhaps required to make tradeoffs in their other behaviors, such as courtship and mating, to minimize energy expenditure. However, prior work indicates rodents have nuanced relationships between ghrelin and mating behavior, as discussed above. Nevertheless, our results demonstrate that male mice with elevated ghrelin de-prioritize courtship behavior by producing fewer vocalizations, suggesting that when ghrelin levels are high, motivation may be shifted away from mating opportunities, potentially in favor of feeding. However, as we did not directly test this hypothesis, future studies should investigate the role of ghrelin in influencing the choice between feeding and mating opportunities.

Our further examination of USV call length and bout length was used to evaluate vocal effort and “performance” (Nicolakis et al., 2020). Despite a significant decrease in the number of USVs produced, we found that when male mice treated with ghrelin vocalized, they called with similar effort to control males, evidenced by similar call length and bout length, between groups. Increases in USV length and complexity have previously been associated with copulatory success (Nicolakis et al., 2020). Similarly, male mice that produce more vocalizations tend to also produce longer and more complex calls (Kono and Kanno, 2026). Consistent with this earlier study, we found a positive relationship between the number of USVs produced and call length, regardless of treatment. Paired with our results respective to call length, bout length, and bout number, these findings suggest the motivation males have to make more effortful vocalizations is somewhat independent of their motivation to vocalize.

Ghrelin is not the only hormone associated with feeding and energy balance that plays an important role in modulating social interactions. Indeed, others have noted the striking overlap between hormonal and neural circuit mechanisms of feeding and social behavior (Fischer and O’Connell, 2017). For example, leptin, a hormonal signal of high energy balance, increases the amplitude of social signals in electric fish (Sinnett and Markham, 2015) and promotes increased display effort in Neotropical singing mice (Giglio and Phelps, 2020; Tripp et al., 2026). On the other hand, neurons in the arcuate nucleus expressing agouti-related peptide–which detect ghrelin and leptin, signal hunger, and promote feeding–inhibit maternal behaviors in female mice (Alcantara et al., 2025; Li and Yan, 2018). While this field of research is growing, it is clear that expression of social behavior can be profoundly influenced by an individual’s energy balance, particularly when behaviors are energetically costly or have clear trade-offs with foraging and feeding such as courtship display or parental care.

Finally, there are two important limitations to our study that should be noted. First, our experiment was conducted in the light phase, in which mice are less active (Pernold et al., 2023). Additionally, all of our female stimulus animals were ovariectomized and were not hormonally primed to be sexually receptive. Both of these factors may have decreased general levels of activity and sexual or social motivation in our subjects. It is plausible that more highly motivated subjects would have displayed greater behavioral differences due to hormone treatment, though also possible that reduced baseline motivation revealed effects of ghrelin we would not have seen in very highly motivated subjects. Nevertheless, our results highlight the particular importance of ghrelin signaling for vocalization compared to other courtship interactions. Future studies may elucidate the interaction between ghrelin signaling and circadian rhythm or partner receptivity in regulating male sexual behavior.

In conclusion, we found that ghrelin reduced the rate of USVs made by male mice toward females but did not significantly impact other social or courtship behaviors. While hunger may influence social behaviors more generally, our results indicate that ghrelin signaling is an especially important regulator of vocal behavior. Overall, these results highlight the importance of ghrelin, a hormone typically associated with signaling hunger, in regulating courtship investment. Further research should broaden our understanding of the role of ghrelin and other hormonal signals of energy balance in regulating social behavior by testing their effects on male courtship interactions with sexually receptive females and in intrasexual interactions as well as identifying circuit mechanisms that link hormonal signals of energy balance with social and motivational systems.

## Acknowledgements

We thank Robyn Durand and staff for assistance with animal care, Sarah Meerts and Dianne Rodman for guidance on surgical procedures, Katherine Tschida for advice on behavioral experiments, and John Paul Janik and Yasmine Tesema for assistance with behavioral observations. This work was funded by Carleton College.

## Author Contributions

J.A.T. conceived the study; R.G.B, L.T.B., and J.A.T. performed the experiments, R.G.B. and L.T.B. analyzed behavioral data, M.O.J. and J.A.T. conducted statistical analyses. All authors contributed writing and revising the manuscript and approved the final submission.

## References

Abrahams, M.V., 1993. The trade-off between foraging and courting in male guppies. Animal Behaviour 45, 673–681. 10.1006/anbe.1993.1082

Alcantara, I.C., Li, C., Gao, C., Rodriguez González, S., Mickelsen, L.E., Papas, B.N., Goldschmidt, A.I., Cohen, I.M., Mazzone, C.M., de Araujo Salgado, I., Piñol, R.A., Xiao, C., Karolczak, E.O., Li, J.-L., Cui, G., Reitman, M.L., Krashes, M.J., 2025. A hypothalamic circuit that modulates feeding and parenting behaviours. Nature 645, 981–990. 10.1038/s41586-025-09268-5

Babaei-Balderlou, F., Khazali, H., 2016. Effects of ghrelin on sexual behavior and luteinizing hormone beta-subunit gene expression in male rats. J Reprod Infertil 17, 88–96.

Ball, G.F., Balthazart, J., 2008. How useful is the appetitive and consummatory distinction for our understanding of the neuroendocrine control of sexual behavior? Horm Behav 53, 307–318. 10.1016/j.yhbeh.2007.09.023

Belén Poretti, M., Bianconi, S., Luque, E., Martini, A.C., Vincenti, L., Cantarelli, V., Torres, P., Ponzio, M., Schiöth, H.B., Carlini, V.P., 2023. Role of the hypothalamus in ghrelin effects on reproduction: sperm function and sexual behavior in male mice. Reproduction 165, 123–134. 10.1530/REP-22-0098

Bertoldi, M.L., Luque, E.M., Carlini, V.P., Vincenti, L.M., Stutz, G., Santillán, M.E., Ruiz, R.D., Cuneo, M.F. de, Martini, A.C., 2011. Inhibitory effects of ghrelin on sexual behavior: role of the peptide in the receptivity reduction induced by food restriction in mice. Horm Metab Res 43, 494–499. 10.1055/s-0031-1277228

Castellucci, G.A., Calbick, D., McCormick, D., 2018. The temporal organization of mouse ultrasonic vocalizations. PLoS ONE 13, e0199929. 10.1371/journal.pone.0199929

Chaverri, G., Sandoval-Herrera, N.I., Iturralde-Pólit, P., Romero-Vásquez, A., Chaves-Ramírez, S., Sagot, M., 2021. The energetics of social signaling during roost location in Spix’s disc-winged bats. Journal of Experimental Biology 224, jeb238279. 10.1242/jeb.238279

Coffey, K.R., Marx, R.E., Neumaier, J.F., 2019. DeepSqueak: a deep learning-based system for detection and analysis of ultrasonic vocalizations. Neuropsychopharmacol. 44, 859–868. 10.1038/s41386-018-0303-6

Collier, K., Parsons, S., Czenze, Z.J., 2022. Thermal energetics of male courtship song in a lek-breeding bat. Behav Ecol Sociobiol 76, 36. 10.1007/s00265-022-03141-5

Cummings, D.E., Purnell, J.Q., Frayo, R.S., Schmidova, K., Wisse, B.E., Weigle, D.S., 2001. A preprandial rise in plasma ghrelin levels suggests a role in meal initiation in humans. Diabetes 50, 1714–1719. 10.2337/diabetes.50.8.1714

Egecioglu, E., Prieto-Garcia, L., Studer, E., Westberg, L., Jerlhag, E., 2016. The role of ghrelin signalling for sexual behaviour in male mice. Addict Biol 21, 348–359. 10.1111/adb.12202

Egnor, S.R., Seagraves, K.M., 2016. The contribution of ultrasonic vocalizations to mouse courtship. Current Opinion in Neurobiology, Neurobiology of sex 38, 1–5. 10.1016/j.conb.2015.12.009

Engel, J.A., Jerlhag, E., 2014. Role of appetite-regulating peptides in the pathophysiology of addiction: implications for pharmacotherapy. CNS Drugs 28, 875–86.

Fischer, E.K., O’Connell, L.A., 2017. Modification of feeding circuits in the evolution of social behavior. The Journal of Experimental Biology 220, 92–102. 10.1242/jeb.143859

Friard, O., Gamba, M., 2016. BORIS: a free, versatile open-source event-logging software for video/audio coding and live observations. Methods in Ecology and Evolution 7, 1325–1330. 10.1111/2041-210X.12584

Giglio, E.M., Phelps, S.M., 2020. Leptin regulates song effort in Neotropical singing mice (*Scotinomys teguina*). Animal Behaviour 167, 209–219. 10.1016/j.anbehav.2020.06.022

Hammond, T.J., Bailey, W.J., 2003. Eavesdropping and defensive auditory masking in an australian bushcricket, *Caedicia* (Phaneropterinae: Tettigoniidae: Orthoptera). Behaviour 140, 79–95. http://www.jstor.org/stable/4536012

Hyland, L., Rosenbaum, S., Edwards, A., Palacios, D., Graham, M.D., Pfaus, J.G., Woodside, B., Abizaid, A., 2018. Central ghrelin receptor stimulation modulates sex motivation in male rats in a site dependent manner. Hormones and Behavior 97, 56–66. 10.1016/j.yhbeh.2017.10.012

Ilany, A., Barocas, A., Kam, M., Ilany, T., Geffen, E., 2013. The energy cost of singing in wild rock hyrax males: Evidence for an index signal. Animal Behaviour, Special Issue: Behavioural Plasticity and Evolution 85, 995–1001. 10.1016/j.anbehav.2013.02.023

Kanno, K., Kikusui, T., 2018. Effect of sociosexual experience and aging on number of courtship ultrasonic vocalizations in male mice. jzoo 35, 208–214. 10.2108/zs170175

Keen-Rhinehart, E., Bartness, T.J., 2005. Peripheral ghrelin injections stimulate food intake, foraging, and food hoarding in Siberian hamsters. American Journal of Physiology-Regulatory, Integrative and Comparative Physiology 288, R716–R722. 10.1152/ajpregu.00705.2004

Kim, T.W., Sakamoto, K., Henmi, Y., Choe, J.C., 2008. To court or not to court: reproductive decisions by male fiddler crabs in response to fluctuating food availability. Behav Ecol Sociobiol 62, 1139–1147. 10.1007/s00265-007-0542-8

Klaus, T., Wernisch, B., Zala, S.M., Penn, D.J., 2025. Courtship vocalizations of wild house mice show highly dynamic changes and correlate with male copulatory success. Animal Behaviour 220, 123024. 10.1016/j.anbehav.2024.11.002

Kono, T., Kanno, K., 2026. Linking quantity and acoustic properties of courtship vocalizations in male mice. Behavioural Brain Research 511, 116279. 10.1016/j.bbr.2026.116279

Korbonits, M., Goldstone, A.P., Gueorguiev, M., Grossman, A.B., 2004. Ghrelin—a hormone with multiple functions. Frontiers in Neuroendocrinology 25, 27–68. 10.1016/j.yfrne.2004.03.002

Li, L., 2026. mouse cage. 10.5281/zenodo.5496331

Li, X., Yan, J., 2018. Agrp neurons project to the medial preoptic area and modulate maternal nest-building. Journal of Neuroscience 39, 456–471.

Marconi, M.A., Nicolakis, D., Abbasi, R., Penn, D.J., Zala, S.M., 2020. Ultrasonic courtship vocalizations of male house mice contain distinct individual signatures. Animal Behaviour 169, 169–197. 10.1016/j.anbehav.2020.09.006

Mifune, H., Tajiri, Y., Sakai, Y., Kawahara, Y., Hara, K., Sato, T., Nishi, Y., Nishi, A., Mitsuzono, R., Kakuma, T., Kojima, M., 2020. Voluntary exercise is motivated by ghrelin, possibly related to the central reward circuit. Journal of Endocrinology 244, 123–132. 10.1530/JOE-19-0213

Mitoyen, C., Quigley, C., Fusani, L., 2019. Evolution and function of multimodal courtship displays. Ethology 125, 503–515. 10.1111/eth.12882

Mougeot, F., Bretagnolle, V., 2000. Predation as a cost of sexual communication in nocturnal seabirds: an experimental approach using acoustic signals. Animal Behaviour 60, 647–656. 10.1006/anbe.2000.1491

NIAID Visual & Medical Arts, 2024. Syringe. https://doi.org/bioart.niaid.nih.gov/bioart/505

Nicolakis, D., Marconi, M.A., Zala, S.M., Penn, D.J., 2020. Ultrasonic vocalizations in house mice depend upon genetic relatedness of mating partners and correlate with subsequent reproductive success. Front Zool 17, 10. 10.1186/s12983-020-00353-1

Noren, D.P., Holt, M.M., Dunkin, R.C., Williams, T.M., 2013. The metabolic cost of communicative sound production in bottlenose dolphins (*Tursiops truncatus*). Journal of Experimental Biology jeb.083212. 10.1242/jeb.083212

Ophir, A.G., Schrader, S.B., Gillooly, J.F., 2010. Energetic cost of calling: general constraints and species-specific differences. J of Evolutionary Biology 23, 1564– 1569. 10.1111/j.1420-9101.2010.02005.x

Pernold, K., Rullman, E., Ulfhake, B., 2023. Bouts of rest and physical activity in C57BL/6J mice. PLoS ONE 18, e0280416. 10.1371/journal.pone.0280416

Prieto-Garcia, L., Egecioglu, E., Studer, E., Westberg, L., Jerlhag, E., 2015. Ghrelin and GHS-R1A signaling within the ventral and laterodorsal tegmental area regulate sexual behavior in sexually naïve male mice. Psychoneuroendocrinology 62, 392–402. 10.1016/j.psyneuen.2015.09.009

Shah, S.N., Nyby, J.G., 2010. Ghrelin’s quick inhibition of androgen-dependent behaviors of male house mice (*Mus musculus*). Hormones and Behavior 57, 291–296. 10.1016/j.yhbeh.2009.12.010

Sinnett, P.M., Markham, M.R., 2015. Food deprivation reduces and leptin increases the amplitude of an active sensory and communication signal in a weakly electric fish. Hormones and Behavior 71, 31–40. 10.1016/j.yhbeh.2015.03.010

Sovetkina, A., Nadir, R., Fung, J.N.M., Nadjarpour, A., Beddoe, B., 2020. The physiological role of ghrelin in the regulation of energy and glucose homeostasis. Cureus 12, e7941. 10.7759/cureus.7941

Tena-Sempere, M., 2007. Roles of Ghrelin and Leptin in the Control of Reproductive Function. Neuroendocrinology 86, 229–241. 10.1159/000108410

Tezenas Du Montcel, C., Cao, J., Mattioni, J., Hamelin, H., Lebrun, N., Ramoz, N., Gorwood, P., Tolle, V., Viltart, O., 2023. Chronic food restriction in mice and increased systemic ghrelin induce preference for running wheel activity. Psychoneuroendocrinology 155, 106311. 10.1016/j.psyneuen.2023.106311

Tripp, J.A., Raghuraman, K., Bhalla, R.V., Phelps, S.M., 2026. Leptin promotes social context-specific increase in advertisement song effort of male Alston’s singing mice. Hormones and Behavior 178, 105874. 10.1016/j.yhbeh.2025.105874

Tritos, N.A., Kokkotou, E.G., 2006. The physiology and potential clinical applications of ghrelin, a novel peptide hormone. Mayo Clinic Proceedings 81, 653–60.

Tuttle, M.D., Ryan, M.J., 1981. Bat predation and the evolution of frog vocalizations in the neotropics. Science 214, 677–678.

Yin, Y., Li, Y., Zhang, W., 2014. The growth hormone secretagogue receptor: its intracellular signaling and regulation. Int J Mol Sci 15, 4837–4855. 10.3390/ijms15034837

